# Identification of novel sex determination loci in Japanese weedy melon

**DOI:** 10.1101/2023.01.20.524881

**Authors:** Akito Nashiki, Hiroki Matsuo, Kota Takano, Fauziatul Fitriyah, Sachiko Isobe, Kenta Shirasawa, Yosuke Yoshioka

## Abstract

Sex expression contributes to fruit quality and yield in the Cucurbitaceae. In melon, orchestrated regulation by sex determination genes explains the mechanism of sex expression, resulting in a great variety of sexual morphologies. In this study, we examined the Japanese weedy melon UT1, which does not follow the reported model of sex expression. We conducted QTL analysis using F_2_ plants for flower sex on the main stem and the lateral branch and mapped a “femaleness” locus on Chr. 3 (*Fem3.1*) and a “type of flower femaleness” (female or bisexual) locus on Chr. 8 (*tff8.1*). *Fem3.1* included the known sex determination gene *CmACS11*. Sequence comparison of *CmACS11* between parental lines revealed three nonsynonymous SNPs. A CAPS marker developed from one of the SNPs was closely linked to femaleness in two F_2_ populations with different genetic backgrounds. The femaleness allele was dominant in F_1_ lines from crosses between UT1 and diverse cultivars and breeding lines. This study suggests that the identity of *tff8.1* is *CmCPR5*, a recently reported bisexual flower control gene. We found that the Japanese weedy melon UT1 does not follow the conventional sex expression model because of the interaction of the loci *Fem3.1* and *tff8.1* with the previously reported sex determination genes. The results of this study provide new insights into the molecular mechanisms of sex determination in melons and considerations for the application of femaleness in melon breeding.

**Key message:** Japanese weedy melon exhibits unique sex expression with interactions between previously reported sex determination genes and two novel loci.

## Introduction

Sex determination plays a pivotal role in the survival strategies of plants and animals. Hermaphrodite (only bisexual flowers) have pistils and stamens and are thought to be the ancestral sexual form and are the most common one, whereas very few plant species are dioecious (male and female plants) or monoecious (male and female flowers on the same plant) (Ainsworth 2000; Barrett 2002; Renner 2014). Although a two-locus model has been proposed for dioecious plants (Charlesworth and Charlesworth, 1978; Renner, 2016), recent genetic studies have revealed diverse sex determination mechanisms in persimmon (*Diospyros* spp.) (Akagi et al. 2014, 2020; Masuda et al. 2019), kiwifruit (*Actinidia* spp.) (Akagi et al. 2018, 2019), garden asparagus (*Asparagus officinalis* L.) (Harkess et al. 2017, 2020; Murase et al. 2017; Tsugama et al. 2017), and poplar (*Populus* spp.) (Müller et al. 2020). In some monoecious plants such as maize and some species in the Cucurbitaceae, the sex determination mechanism depends on the endogenous production or exogenous application of plant hormones (Hartwig et al. 2011; Chen et al. 2016). In the Cucurbitaceae, ethylene is crucial in sex determination (Byers et al. 1972). In cucumber (*Cucumis sativus* L.) and melon (*Cucumis melo* L.), male, female, and bisexual flowers can occur on the same plant. Male and female flowers differentiated from bisexual flowers as a result of a selective arrest in the development of the stamen or carpel primordium (Kater et al. 2001; Hao et al. 2003; Bai et al. 2004). Thus, cucumber and melon have a great variety of sexual morphologies, including not only monoecious but also andromonoecious (male and hermaphrodite flowers on the same plant), hermaphrodite, gynoecious (only female flowers), and androecious (only male flowers). Elucidation of the network of hormones and genes that control sex expression in cucurbits will not only provide important knowledge for evolutionary research on plant sexuality, but will also have critical applications in plant breeding and crop production, because sex determination is essential for maintaining yield, improving fruit quality, and planning breeding strategies in cucurbits (Grumet and Taft, 2012).

At least three independent significant genes have explained the mechanism of sex determination in melon, *CmACS-7* (locus *m*; Boualem et al. 2008) and *CmACS11* (locus *a*; Boualem et al. 2015) encode for 1-aminocyclopropane-1-carboxylate (ACC) synthase, which catalyzes the rate-limiting step in ethylene biosynthesis, and *CmWIP1* (locus *g*; Martin et al. 2009), a C2H2 zinc transcription factor. Allelic variations of these genes result in different sex expression: hermaphrodite (*--ggmm*), andromonoecious (*A-G-mm*), gynoecious (*--ggM-*), monoecious (*A-GGM-*), and androecious (*aaGG--*) (Boualem et al. 2015). Thus, plants homozygous for the recessive allele (*g*) of *CmWIP1* have only female or bisexual flowers. *CmACS11* represses the expression of the male-promoting gene *CmWIP1* in lateral branches, but allows *CmWIP1* expression in the main stem to control the development and coexistence of male and female flowers in monoecious species (Boualem et al. 2015).

However, a Japanese weedy melon accession (UT1) collected from an island in the Seto Inland Sea has the *GGMM* genotype and is expected to be monoecious, but it is hermaphrodite. Interestingly, many wild melon accessions reportedly harbor the *CmACS-7* haplotype (*M*) and should be monoecious, but are andromonoecious (Zhang et al. 2019). In oriental melon, GWAS suggested two novel SNPs associated with sex expression, on chromosomes (Chr.) 1 and 8 (Kishor et al. 2021). A novel bisexual flower control gene has been identified on Chr. 8 of a hermaphroditic melon germplasm (Wang et al. 2022), and a novel sex determination gene, *CmCRC*, has been identified on Chr. 10 (Zhang et al., 2022). Thus, the study of the genetic mechanism of melon sex determination has much potential, and novel genes may still be discovered in melon.

The purpose of this study was to clarify the inheritance of sex expression in UT1 using a biparental mapping population, and to develop a DNA marker linked to a novel gene related to femaleness on the main stem.

## Materials and Methods

### Plant materials

UT1 is a hermaphrodite weedy melon (*Cucumis melo* L.) accession that belongs to the horticultural group agrestis; it bears bisexual flowers on the main stem and lateral branches (Fig. 1). ‘Earl’s Favourite Harukei 3’ (EF) is an andromonoecious muskmelon cultivar (cantalupensis-reticulatus group); it bears male flowers on the main stem and bisexual flowers on the lateral branches. We used F_2_ plants derived from F_1_ [EF (♀) x UT1 (♂)] (136 plants in September-December 2015, 140 in March-June 2017, and 120 in August–November 2020), and 124 F_2_ plants from F_1_ [UT1 (♀) x EF (♂)] in February–May 2021. To assess the availability of a CAPS marker for femaleness (phenotype with female and bisexual flowers) on the main stem, we used 60 F_2_ plants derived from a cross between UT1 and the non-netted andromonoecious fixed line HOF (inodorous group). To evaluate the effect of a femaleness allele in diverse melon cultivars, we used 3 plants of each F_1_ progeny from crosses between UT1 and andromonoecious cultivars or fixed lines from the inodorous group (HOF, non-netted fixed line SIF, Hami melon cultivar ‘Hamiuri’, and ‘Piel de Sapo’), from the cantalupensis–reticulatus group (EF, ‘Vedrantais’, ‘Charentais’, and from the momordica group (Japanese landrace fixed line).

**Fig. 1.**
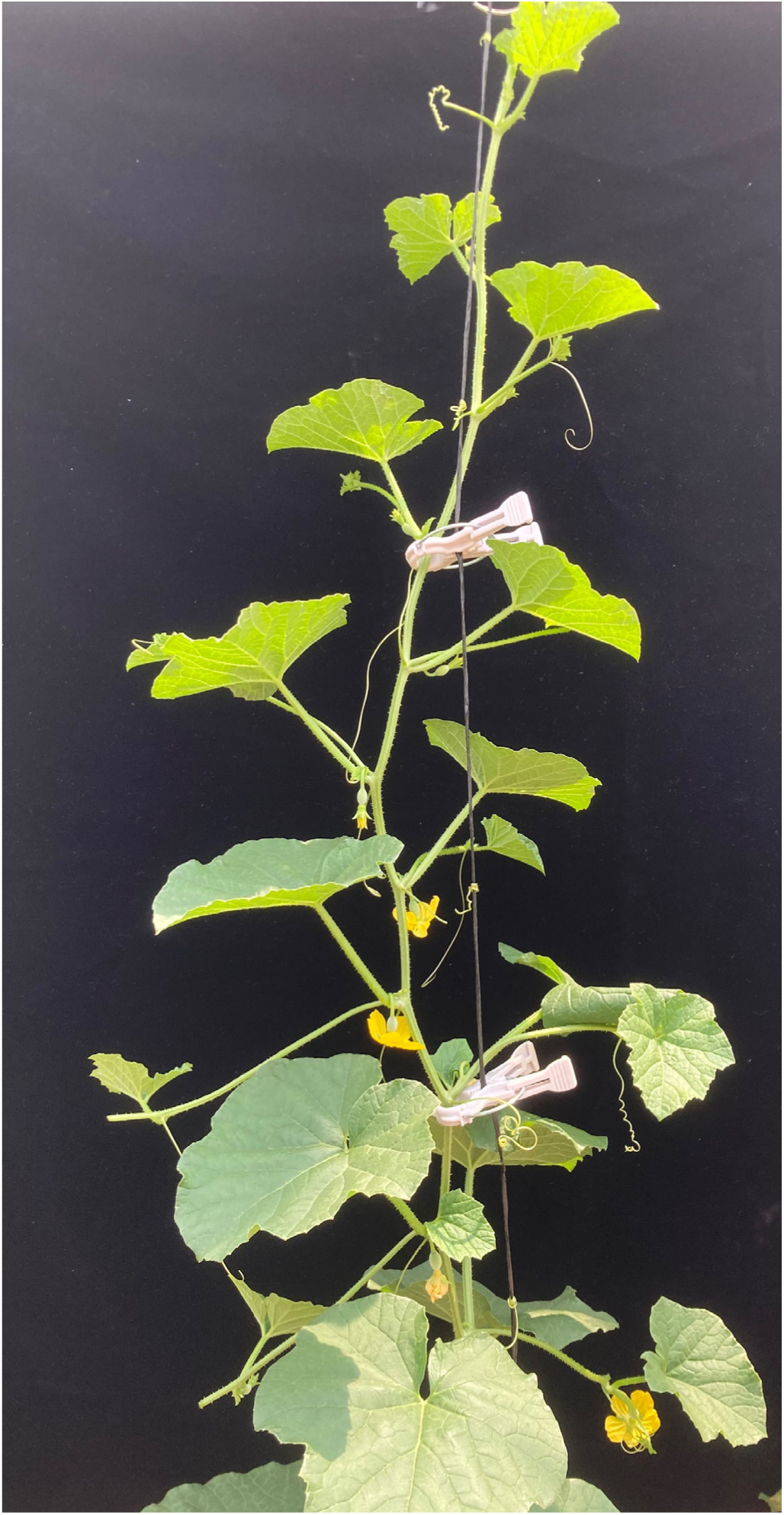
Phenotype of hermaphrodite Japanese weedy melon UT1

### Phenotyping of sex expression

All plants were sown in 6-cm-diameter plastic pots filled with moist culture soil (Nihon Hiryo Co., Ltd., Tokyo, Japan). On day 14, seedlings were transplanted into a rectangular planter box (30 L) filled with wet peat moss soil (Hokkaido Peat Moss Inc., Konosu, Japan) in a plastic greenhouse at the University of Tsukuba, Japan. In 2015, the sex types (female, bisexual, male) of flowers at 8–21 nodes on the main stem were recorded, and the phenotype of each plant was categorized into femaleness (female and bisexual flowers), maleness (male flowers), or mixed (female, male, and bisexual flowers). In subsequent cultivations, the sex types of fully opened flowers at 6–21 nodes on the main stem and the first nodes of lateral branches arising from 6–21 nodes of the main stem were recorded. Sex expression of each plant was then determined on the basis of the sex of all flowers. Two flower sex-related traits—femaleness on the main stem and type of flower femaleness (female or bisexual)—were then used in genetic analysis.

### Genotyping of sex determination genes

Genomic DNA was isolated from young leaves of the parental lines and F_1_ and F_2_ populations using the DNeasy Plant Kit 96 (Qiagen, Hilden, Germany). *CmACS-7* and *CmWIP1* were genotyped according to Boualem et al. (2008) and Chen et al. (2016). Amplification was carried out in 10 μL containing 2 μL of total DNA (5 ng/μL), 5 μL of 2x Ampdirect Plus (Shimadzu, Kyoto, Japan). 0.1 μL of Blend Taq Plus (2.5 U/μL) (Toyobo, Osaka, Japan), and 10 μM of forward and reverse primers (Table S1). For *CmWIP1* markers, the amplification products were electrophoresed in 2% agarose gel in 1x TAE buffer. For the *CmACS-7* CAPS marker, the amplification products were digested with *Alu*I (Takara Bio, Shiga. Japan) and separated in 2% agarose gel in 1x TAE buffer.

### ddRAD-Seq analysis

Genomic DNA was isolated from parental lines and F_2_ populations using the DNeasy Plant Kit 96 (Qiagen) and diluted to a final concentration of 25 ng/μL. Library construction and sequencing analysis were performed as described by Shirasawa et al. (2016) with minor modifications. The ddRAD-Seq libraries were constructed with two combinations of restriction enzymes, *Pst*I and *Msp*I (Thermo Fisher Scientific, Waltham, MA, USA). Digested DNA was ligated to adapters using T4 DNA ligase (Takara Bio Inc.) and purified using Agencourt AMPure XP cleanup reagent (Beckman Coulter, Brea, CA, USA) to eliminate short (<300 bp) DNA fragments. Purified DNA was amplified by PCR with indexed primers. PCR fragments were purified using the QIAquick PCR Purification Kit (Qiagen), and fragments (300–1000 bp) were separated by electrophoresis in 2.0% agarose gel (Kanto Chemical Co., Inc., Tokyo, Japan) and extracted with a MinElute Gel Extraction Kit (Qiagen). The libraries were sequenced on a HiSeq 4000 (Illumina Inc., San Diego, CA, USA) in 100-bp paired-end reads. The obtained ddRAD-Seq reads were mapped onto the *C. melo* reference genome, Melon_genome_v3.5.1 (http://cucurbitgenomics.org/organism/3) in Bowtie 2 (Langmead and Salzberg 2012). Variant calling was performed by bcftools 0.1.19 mpileup in SAMtools (Li et al. 2009). Filtering of variants by quality was performed in vcftools.

### Linkage map construction and QTL analysis

Two genetic linkage maps were constructed separately for F_2_ populations derived from reciprocal F_1_s; the maps were based on the SNPs obtained by ddRAD-seq analysis in R/ASMap (Taylor and Butler, 2017) and R/qtl (Arends et al. 2010). The SNP markers were selected by removing low-quality loci (>80% of F_2_ plants miss values at the locus; mapped to Chr. 0). F_2_ plants harboring >80% of all SNP markers were selected. Then *p*-values were calculated to test for deviations of genotype frequencies at each locus from the expected ratio of 1:2:1, and after Bonferroni correction for multiple testing, markers that showed significant distortion at the 5% level were removed. Genotype error rate and genotype error logarithm of the odds (LOD) scores were calculated and SNP genotypes with high genotype-error LOD score (>0) were treated as missing data. Genetic distance, the order of SNP markers, and the imputation of missing genotypes were calculated using the function mstmap in R/ASMap and the minimum-spanning-tree (MST) algorithm. The Kosambi mapping function was applied during the calculation of genetic map distance. QTL analysis of two flower sex–related traits was performed by non-parametric interval mapping under a single-QTL model in R/qtl. We also calculated the 1.5-LOD support intervals for the location of the identified QTLs.

### Prediction of candidate genes, sequencing, and CAPS analysis

Candidate genes within the 1.5-LOD intervals of QTLs were extracted from the Cucurbit Genomics Database (CuGenDB, http://cucurbitgenomics.org/), and 24 primers were designed for sequencing of a candidate gene for femaleness on the main stem and CAPS marker analysis (Table S1). Amplification was carried out in 10 μL containing 2 μL of total DNA (5 ng/μL), 5 μl of 2x Ampdirect Plus, 0.1 μL of Blend Taq Plus (2.5 U/μL), and 10 μM each forward and reverse primers. The thermal profile was as follows: 95 °C for 2 min; 35 cycles at 95 °C for 30 s, 55 °C for 30 s, and 72 °C for 1 min; and a final extension at 72 °C for 5 min. The PCR products were Sanger-sequenced by a commercial provider (Fasmac Co., Ltd., Kanagawa, Japan). We used *CmACS11* sequence polymorphisms with parental lines to develop a CAPS marker, which permitted *Alu*I digestion from the 5’ ends of both EF alleles but not of UT1 alleles (Table S1). The digested PCR products were electrophoresed in 2.0% agarose gel in 1x TAE buffer. The CAPS marker was applied to the two F_2_ populations in 2020 and 2021, and the relationships between the genotype and phenotype were examined.

## Results

### Inheritance of sex expression

Sex expression in the parental lines UT1 (Fig. 1; only bisexual flowers) and EF (male flowers on the main stem and bisexual flowers on lateral branches) was consistent throughout the three seasons, indicating stability of sex expression in different environments. The F_1_ plants had male, female, and bisexual flowers on the main stem, and female or bisexual flowers on lateral branches. In 2015, sex type on the main stem of the F_2_ population segregated, and 65.4% of F_2_ plants had pistil-bearing flowers on the main stem (Table S2). In 2017, 2020, and 2021, hermaphrodite, andromonoecious, monoecious, gynoecious, gynomonoecious, and trimonoecious plants were detected (Table 1). The genotypes of *CmWIP1* (*G*/*g*) and *CmACS-7* (*M*/*m*) of the parental lines were *GGMM* in UT1 and *GGmm* in EF. According to the sex determination model of Boualem et al. (2015), the genotypes *GGMM* and *GGM-* produce a monoecious phenotype, but the UT1 and F_1_ phenotypes were not monoecious. In 2017, 2020, and 2021, the χ^2^ test showed that the segregation ratio of genotypes (*GGMM*, *GGMm*, *GGmm*) followed the expected 1:2:1 ratio. Since *GGMM* and *GGMm* plants are monoecious and *GGmm* plants are hermaphrodites (Boualem et al. 2015), the expected phenotypes were monoecious and hermaphroditic with an expected segregation ratio of 3:1. However, the phenotypes of the F_2_ populations were differed completely from the expected ones.

**Table 1.** Sex expression phenotypes in 2017, 2020, and 2021.

| Sex expression | March–June 2017 | August–November 2020 | February–May 2021 |
| --- | --- | --- | --- |
| Andromonoecious | 58 | 18 | 31 |
| Monoecious | 17 | 5 | 17 |
| Hermaphrodite | 73 | 57 | 37 |
| Gynoecious | 11 | 24 | 29 |
| Gynomonoecious | 0 | 6 | 8 |
| Trimonoecious | 0 | 10 | 2 |

### Linkage map and QTL analysis

The total number of reads from F_2_ plants was 127 863 716 in September–December 2015; 132 233 099 in March–June 2017; 239 488 250 in August–November 2020; and 152 686 656 in February–May 2021; 2549, 2667, 2018, and 2550 SNPs, respectively, were obtained from these F_2_ plants.

Linkage maps were constructed with SNP markers in the F_2_ populations derived from F_1_ between EF (♀) x UT1 (♂) (1666 markers) and between UT1 (♀) x EF (♂) (2258 markers). The maps contained 12 linkage groups spanning 1443.4 cM in the former F_2_ population and 1240.8 cM in the latter one, with an average marker distance of 0.9 cM and 0.6 cM, respectively. We detected one QTL for femaleness on the main stem on Chr. 3 (*Fem3.1*) in all four cultivations and two QTLs for the type of flower femaleness (female or bisexual) on Chr. 2 (*tff2.1*) in two cultivations and on Chr. 8 (*tff8.1*) in three cultivations (Fig. 2, Table 2). The peak LOD values of *Fem3.1* ranged from 12.05 to 21.07, those of *tff2.1* ranged from 4.11 to 4.14, and those of *tff8.1* ranged from 10.54 to 13.73. Plants with *GGMM* or *GGMm* homozygous for the UT1 allele of *tff8.1* had bisexual flowers (Table 3).

**Fig. 2.**
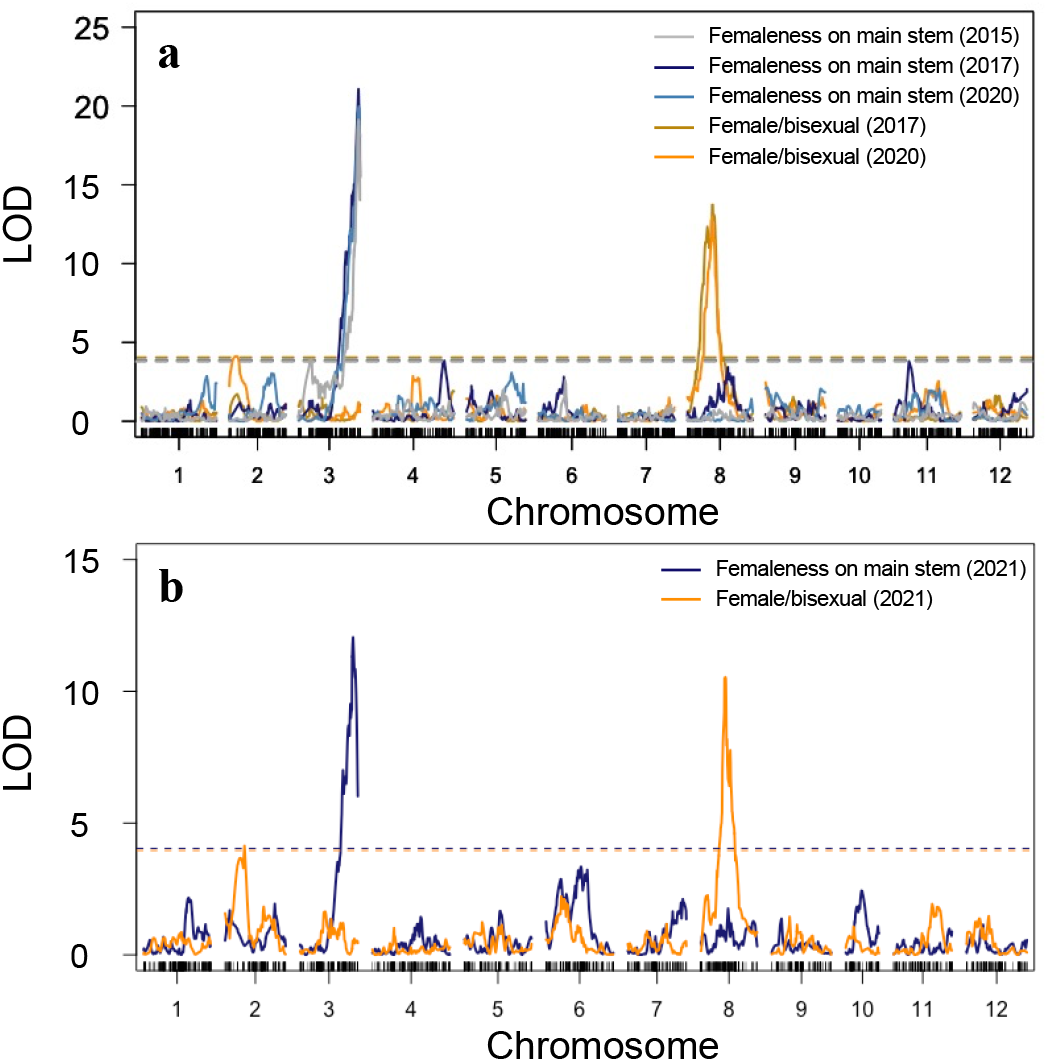
QTL analysis of sex expression. Positions and LOD scores of QTLs associated with two sex-related flower traits—femaleness on the main stem and type of femaleness (female or bisexual flowers)—in F_2_ population in (a) 2015, 2017, and 2020 [EF (♀) × UT1 (♂)] and (b) 2021 [UT1 (♀) × EF (♂)]. Horizontal lines show LOD thresholds at the 0.05 level of significance

**Table 2.** QTLs for sex expression detected by non-parametric single-interval mapping in F_2_ populations from EF x UT1 and UT1 x EF.

| Trait | Chr. | Position (cM) | LOD peak | Estimated range within ‘DHL92’<br>v. 3.5.1 genome | Cultivation |
| --- | --- | --- | --- | --- | --- |
| *Femaleness on main stem* |  |  |  |  |  |
|  | 3 | 116.3 | 19.1 | 28460690–28993081 | September–December 2015 |
|  | 3 | 116.3 | 21.07 | 27857096–28993081 | March–June 2017 |
|  | 3 | 117 | 19.96 | 28460690–28993081 | August–November 2020 |
|  | 3 | 90.7 | 12.05 | 28198809–28993081 | February–May 2021 |
| *Type of femaleness flower (female or bisexual flowers)* |  |  |  |  |  |
|  | 8 | 49 | 13.73 | 4213811–6025370 | March–June 2017 |
|  | 2 | 14 | 4.11 | 597673–2524589 | August–November 2020 |
|  | 8 | 48 | 12.98 | 5081027–6025370 | August–November 2020 |
|  | 8 | 42 | 10.54 | 5214094–6142579 | February–May 2021 |
|  | 2 | 33.3 | 4.14 | 1298407–3048199 | February–May 2021 |

**Table 3.** Relationship between genotype of sex determination genes, allele at *tff8.1*, and sexual type of the population in 2020.

| Genotype<sup>1</sup> | Expected phenotype<sup>2</sup> | Allele at *tff8.1*<sup>3</sup> | Sexual type |  |  |
| --- | --- | --- | --- | --- | --- |
|  |  |  | Female | Bisexual | Both |
| *GGMM* | Female | UT1 | 0 | 8 | 0 |
|  |  | EF | 9 | 0 | 0 |
|  |  | Heterozygous | 3 | 10 | 2 |
| *GGMm* | Female | UT1 | 0 | 9 | 0 |
|  |  | EF | 12 | 2 | 4 |
|  |  | Heterozygous | 2 | 26 | 6 |
| *GGmm* | Hermaphrodite | UT1 | 0 | 6 | 1 |
|  |  | EF | 1 | 3 | 0 |
|  |  | Heterozygous | 0 | 11 | 2 |

<sup>1</sup> G/g: *CmWIP1*; M/m: *CmACS-7*
<sup>2</sup> Expected phenotype based on the genotypes of *CmWIP1* and *CmACS-7*
<sup>3</sup> Based on the SNP marker located on the peak of *tff8.1*

**Table 4.**
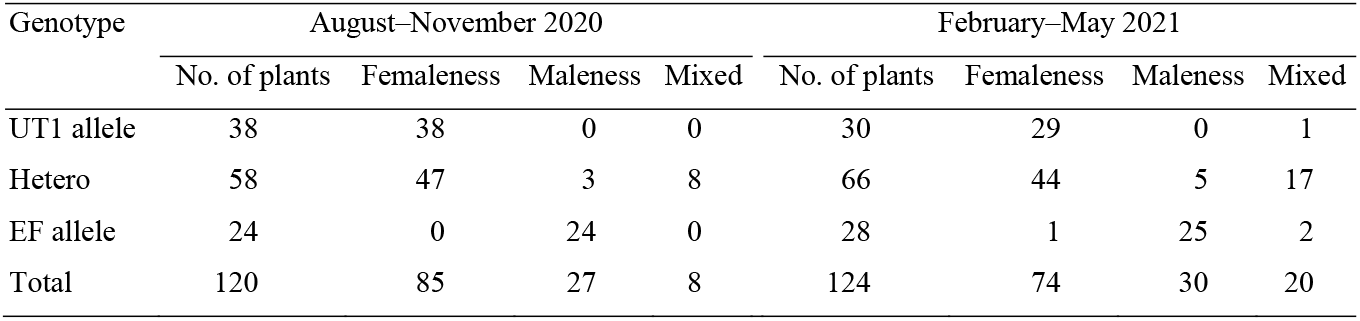
Genotype of the CAPS marker on Chr. 3 and frequency of femaleness or maleness on the main stem in two F_2_ populations.

### Prediction of candidate genes and CAPS marker analysis

Using the melon genome ‘DHL92’ v. 3.5.1 annotation database, we annotated genes within the 1.5-LOD intervals of the QTLs at *tff8.1* (262 genes) and *Fem3.1* (76 genes, including the sex determination gene *CmACS11*; Boualem et al. 2015) (Tables S3, S4).

One of the genes at *tff8.1* was *CmCPR5*, which has recently been reported as a candidate gene controlling bisexual flowers (Wang et al. 2022). Therefore, we sequenced *CmCPR5* and detected polymorphism between the parental lines. The polymorphism of UT1 was consistent with that of the bisexual line Y101 (Wang et al. 2022) (Fig. 3). We further investigated the relationship between the phenotype in 2020 and the genotype of a SNP marker adjacent to *CmCPR5* used in the RAD-seq analysis. We were able to genotype 117 of the 120 plants, and found that the SNP marker S8_5329551, located approximately 30 kbp away from *CmCPR5*, showed strong a relationship with the phenotype (Table S5).

**Fig. 3.**
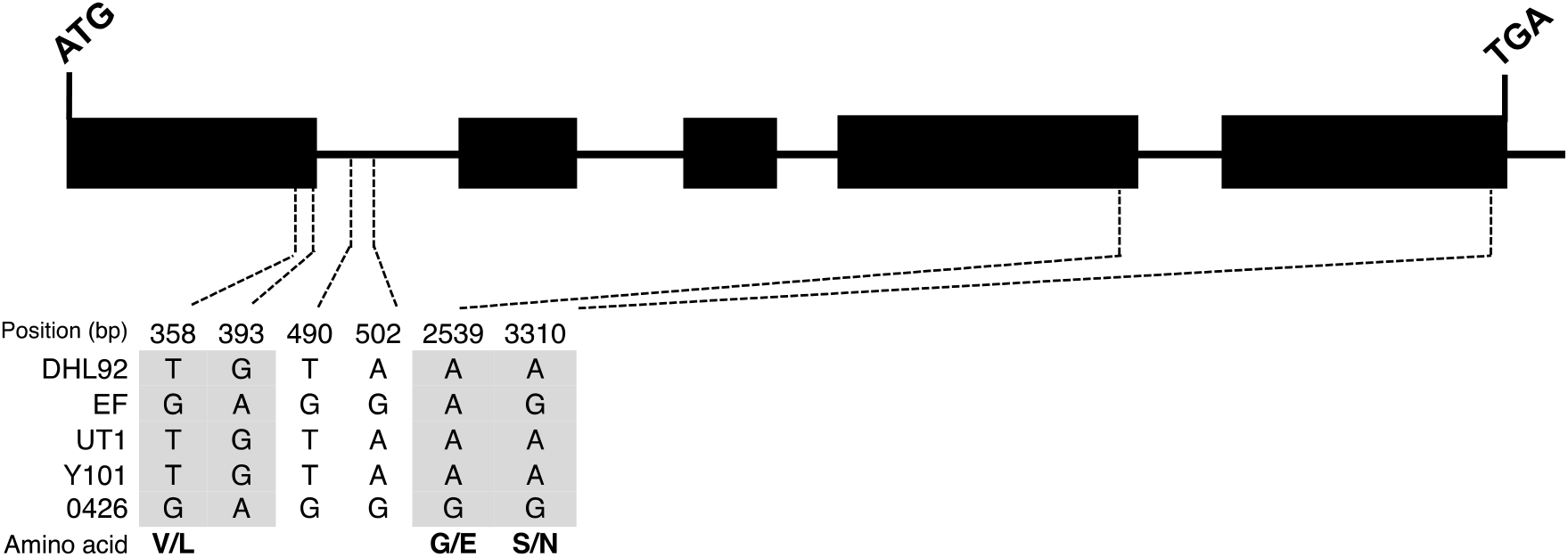
*CmCPR5* sequence variation among ‘DHL92’ (reference genome), EF and UT1 (sequenced in this study), and the hermaphrodite line Y101 and monoecious line 0426 (Wang et al. 2022). Gray shading indicates SNPs in the coding sequence. Amino acid substitutions in ‘DHL92’ or EF relative to UT1 and Y101 are shown in bold; the first amino acid corresponds to ‘DHL92’ or EF and the second one to UT1

We sequenced *CmACS11* in the *Fem3.1* region and detected polymorphisms between the parental lines. We found eight SNPs in the coding region of *CmACS11*, including three non-synonymous substitutions between EF and UT1 (Fig. 4). We developed a CAPS marker according to the SNP at 1050 bp from the start codon in *CmACS11*. CAPS marker analysis showed that the UT1 allele was closely linked to femaleness on the main stem in the F_2_ populations in 2020 and 2021, except in one plant (Table 4). Heterozygotes exhibited femaleness (most of the plants), maleness, or a mixture on the main stem. We also applied CAPS analysis to another F_2_ population, derived from a cross between the non-netted andromonoecious fixed line HOF (♀), whose genotype is *GGmm*, and UT1 (♂), and obtained results similar to those with the F_2_ population between EF and UT1; the UT1 allele was closely linked to femaleness on the main stem (Table S6).

**Fig. 4.**
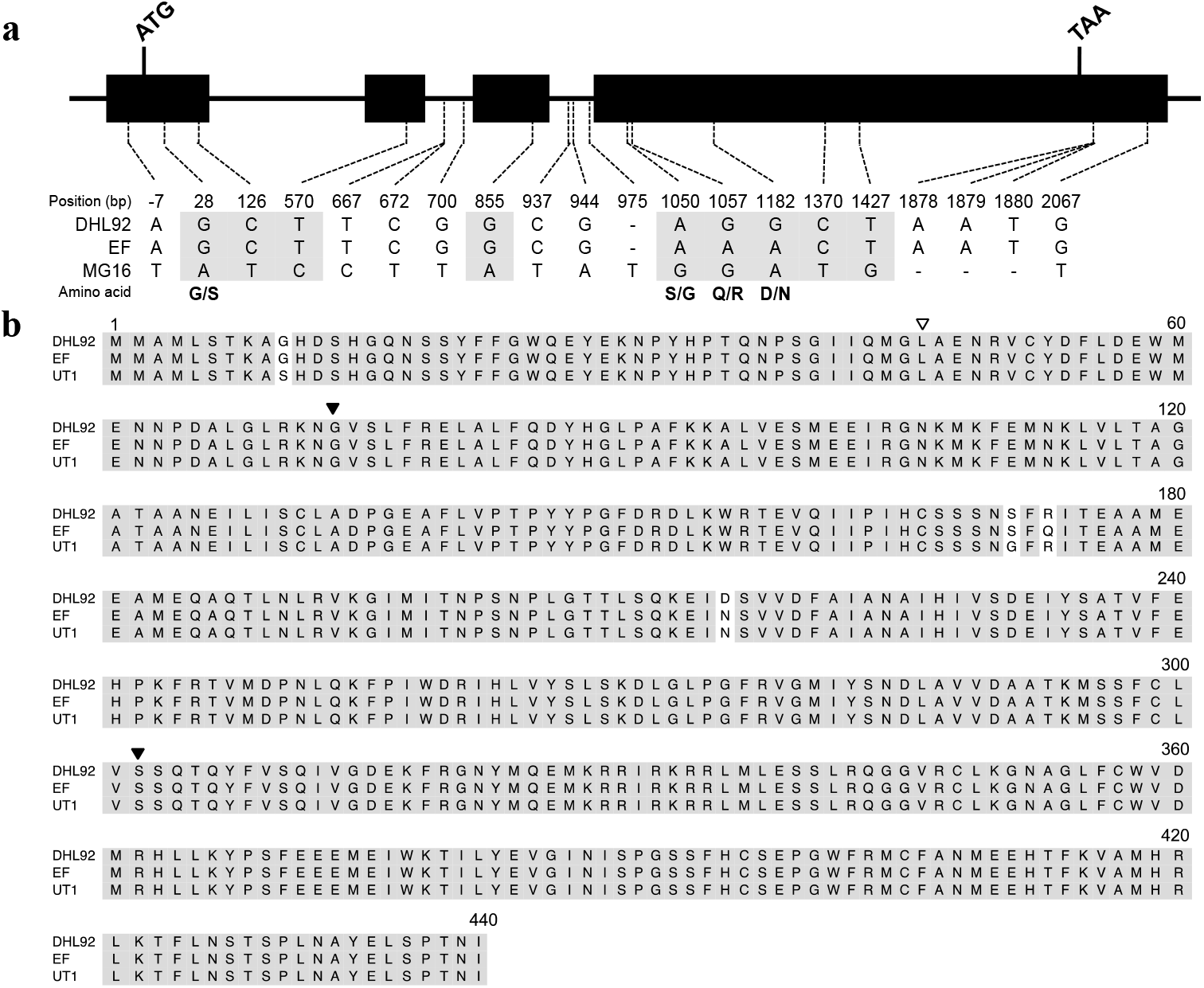
Molecular characterization of the *CmACS11* gene and protein. (a) *CmACS11* sequence variation among ‘DHL92’ (reference genome), EF, and UT1. SNP positions are numbered from the first nucleotide of the start codon. Gray shading indicates SNPs in the coding sequence. Amino acid substitutions in ‘DHL92’ or EF relative to UT1 are shown in bold; the first amino acid corresponds to ‘DHL92’ or EF and the second one to UT1. (b) Amino acid alignment of CmACS11. White arrowhead indicates the reported nonsynonymous substitution among EMS-induced mutations that does not change the monoecious phenotype; black arrowheads indicate nonsynonymous substitutions that change the phenotype from monoecious to androecious (Boualem et al. 2015)

### Sex expression in F_1_s from crosses between UT1 and diverse andromonoecious lines

The genotypes of *CmWIP1* (G/g) and *CmACS-7* (M/m) of all andromonoecious lines were *GGMM*. Thus, those of all F_1_s were *GGMm*; therefore, as mentioned above, the phenotype of all F_1_s was expected to be monoecious. However, all F_1_s showed femaleness on the main stem. On the other hand, flower sex varied among pollen parents. All but two F_1_ lines were hermaphrodites, while F_1_ [UT1 (♀) x EF (♂)] was trimonoecious and F_1_ [UT1 (♀) x SIF (♂)] was gynoecious (Table 5).

**Table 5.**
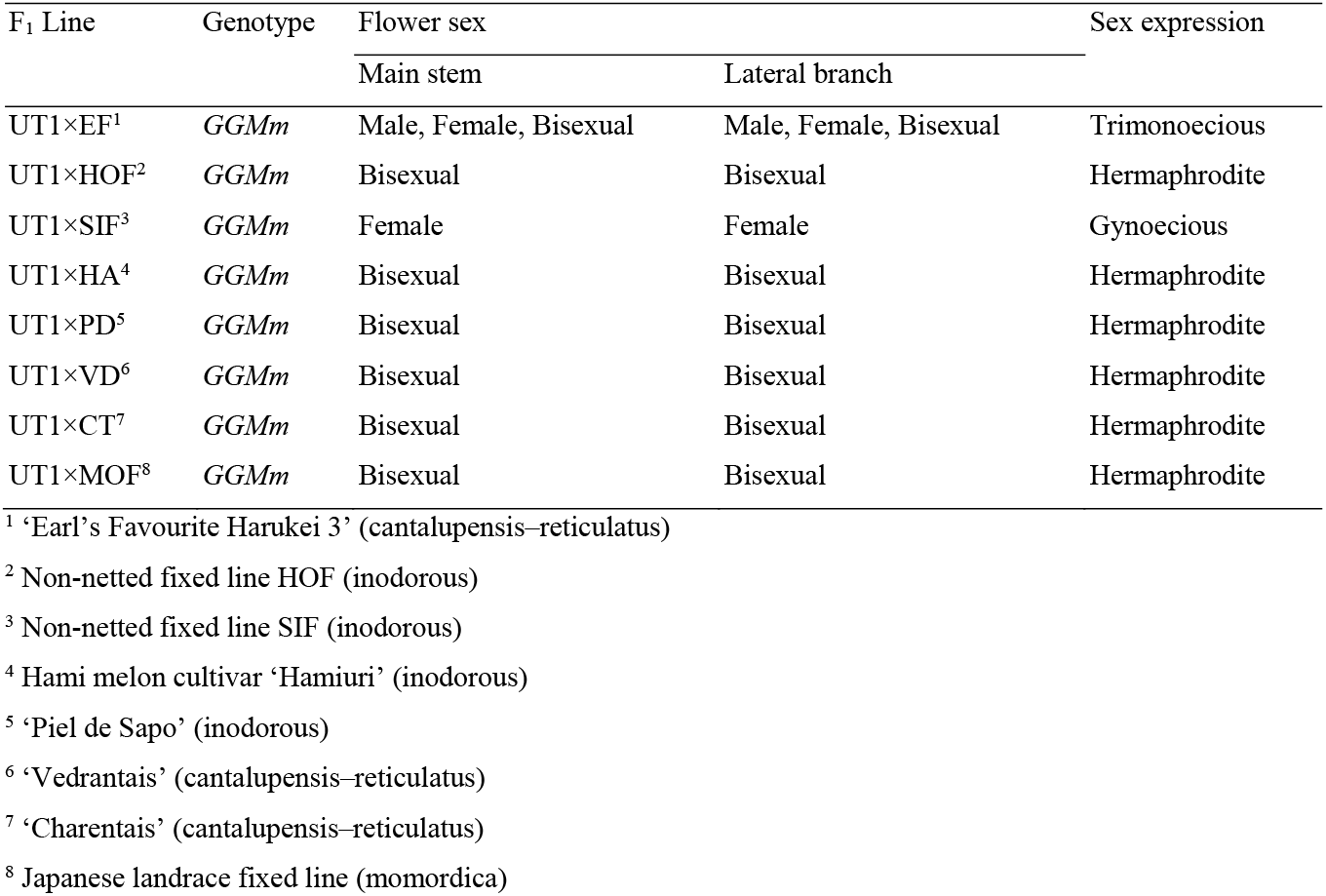
Genotypes and phenotypes of F_1_ lines from crosses between UT1 and andromonoecious lines.

## Discussion

The genetic mechanism of sex expression in melons has hitherto been explained as the orchestrated action of *CmACS11* (Boualem et al. 2015), *CmACS-7* (Boualem et al. 2008), and *CmWIP1* (Martin et al. 2009), which regulate differentiation and development of male and female organs, resulting in the formation of bisexual, female, and male flowers. However, the weedy melon accession UT1 is a hermaphrodite, although its genotype is *GGMM*, which makes plants monoecious according to the sex expression model of Boualem et al. (2015). The sex expression of F_1_ from crosses between UT1 and EF and most of the F_2_ plants differed from that predicted by the model. Our QTL analysis revealed three QTLs for sex expression (Fig. 2), including a QTL region for femaleness on the main stem (*Fem3.1*) that harbored *CmACS11*. We detected three nonsynonymous SNPs between UT1 and EF in a *CmACS11* sequence alignment (Fig. 4) and developed a CAPS marker in *CmACS11*. This marker was closely linked to femaleness on the main stem that was exhibited by plants homozygous for the UT1 allele. Most of heterozygous plants also exhibited femaleness, but several plants exhibited a mixture or maleness (Tables 4, S4). This indicates that the femaleness caused by the UT1 allele of the QTL is dominantly inherited. Melon *CmACS11* and cucumber *CsACS11* are not expressed in the main stem; in melon, expression in lateral branches suppresses *CmWIP1* expression, resulting in female flowers. Loss-of-function mutants of *CmACS11* confer androecy (Boualem et al. 2015). In cucumber, larger copy number of the *CsACS1/G* genes encoding ACC synthase causes femaleness (Li et al. 2020), but the contribution of *CmACS11* to femaleness has not been reported.

We suggest that the causal factor for femaleness might be (1) a novel allele of *CmACS11* derived from UT1, (2) a novel gene closely linked to *CmACS11*, or (3) structural mutations surrounding *CmACS11*. Some heterozygous *CmACS11* plants had male flowers on the main stem. Heterozygous alleles may not be enough for femaleness and are influenced by some other QTLs or environmental factors (Miao et al. 2011; Manzano et al. 2014; Bu et al. 2016; Win et al. 2019). The genetic network mechanism of sex expression in monoecious plants has been studied (Latrasse et al. 2017; Rodriguez-Granados et al. 2022); the femaleness allele detected here may shed light on such a mechanism. Investigating the genomic structure of the region around *CmACS11* using long-read sequencing analysis of UT1 and expression analysis of each reported sex determination gene will provide additional insight.

Of the two QTLs for the type of flower femaleness, that on Chr. 2 (*tff2.1*) could be due to *CmACS-7* because the candidate region included this gene. The F_2_ plants derived from a cross between EF and UT1 that were genotyped *GGMM* or *GGMm* for *CmWIP1* and *CmACS-7* and had no UT1 allele on Chr. 8 (*tff8.1*) bore female flowers (Table 3), in line with the sex determination model. This result supports the identity of *tff2.1* as *CmACS-7*. As to *tff8.1*, genome-wide association (GWAS) analysis in oriental melons reported a significant SNP for sex expression on Chr. 8 (Kishor et al. 2021). The phenotype of melons with the haplotype *M* due to the A57V mutation in *CmACS-7* is monoecious in landraces and cultivars (Boualem et al. 2008); in contrast, most wild-type melons with the same haplotype A57V are andromonoecious and bear bisexual flowers (Zhang et al., 2019). Recently, Wang et al. (2022) reported a candidate gene, *CmCPR5*, for bisexual flowers with haplotype *M* in *CmACS-7*. In our study, homozygosity for the UT1 allele at *tff8.1* (determined by the SNP marker nearest to the LOD peak) and the SNP marker adjacent to *CmCPR5* indicated andromonoecious or hermaphrodite with haplotype *M* in *CmACS-7* (Tables 3, S4). The sequence of UT1 *CmCPR5* matched that of the reported bisexual line Y101 (Wang et al. 2022), suggesting that *CmCPR5* may contribute to the hermaphrodite and andromonoecious phenotypes with *GGMM*. Therefore, *Fem3.1* and *CmCPR5* may influence the sex expression of weedy melon UT1 by interacting with the reported sex determination genes.

To apply the QTLs detected here in practical breeding, it is essential to characterize their inheritance and genetic effect in diverse genetic backgrounds. In this study, all F_1_s from crosses between UT1 and diverse andromonoecious lines with *GGmm* had femaleness on the main stem. On the other hand, even though the genotypes of all F_1_ lines were *GGMm*, their sex expression differed depending on the pollen parents (Table 5). This may be due to the dominant effect of the femaleness allele at *Fem3.1* derived from UT1 and differences in allele-to-allele interactions between UT1 and the respective paternal lines at *tff8.1* (possibly *CmCPR5*). The recessive *g* allele has been used to create gynoecious lines (Cohen et al. 1993), and *CmWIP1* has been identified as the causal gene (Martin et al. 2009). In our study, the femaleness allele located on Chr. 3 had a dominant effect. Therefore, the novel femaleness allele derived from UT1 offers a dominant effect in crosses with lines of diverse genetic backgrounds and would be useful to confer femaleness on the main stem to F_1_ cultivars of any horticultural group. In hybrid seed production, the use of gynoecious melon lines has been proposed because it saves the cost of emasculation and manual pollination and ensures 100% seed purity (Loy et al. 1979; Kenigsbuch and Cohen 1990). However, current gynoecious lines with the *g* allele are unstable at high temperatures, which limit their use in seed production (Kesh and Kaushik 2021). If a gynoecious line with the femaleness allele detected in this study overcomes this problem, it will significantly contribute to efficient seed production.

Sex expression in cucurbits is an important agronomic trait that can improve fruit quality and yield and contribute to more efficient seed production and harvesting systems. In cucumber, the gynoecious type has long been used in practical breeding for the number of female flowers, which is directly related to yield, since immature fruits are harvested (Takahashi and Suge, 1980). In melon, even though a gynoecious cultivar has been developed (Cohen et al. 1993), most modern cultivars are andromonoecious. Andromonoecious plants change the sex of flowers on the main stem (Karch 1970; Papadopoulou et al. 2005): they bear only male flowers initially, but may bear both male and bisexual flowers later. Plants generally bear a bisexual flower on the first node of a lateral branch that grows from the first node of the main stem (Rudich et al. 1969; Little et al. 2007). The use of gynoecious plants can eliminate the waiting period for the lateral branches to elongate and facilitate fruit set on the main stem. It also contributes to higher yields per unit area through pruning lateral branches and high-density cultivation, as they can produce fruit on the main stem. Therefore, femaleness on the main stem has a potential for utilization in melon breeding.

The sex determination model based on *CmACS-7*, *CmWIP1*, and *CmACS11* has hitherto explained all sex expression found in melon (Boualem et al. 2008, 2015; Martin et al. 2009). The model continues to be updated, and the sex expression genes *CmCPR5* on Chr. 8 and *CmCRC* on Chr. 10 have recently been reported (Wang et al. 2022; Zhang et al. 2022). We found that the Japanese weedy melon UT1 does not follow the conventional sex expression model because of the interaction of the loci *Fem3.1* and *tff8.1* with the previously reported sex determination genes. The identity of *tff8.1* may be *CmCPR5*, whereas *Fem3.1* may be a novel QTL and may provide new insights into the sex determination mechanism.

## Acknowledgments

We acknowledge funding for this work from JSPS KAKENHI and JST SPRING. We thank N. Fujishita at Osaka Prefecture University for providing seeds of Japanese weedy melon. We are grateful to N. Nishi and H. Matsumura for their technical assistance.

## Data availability

Datasets generated and analyzed during this study are available from the corresponding author on reasonable request.

---

**Table S1.**
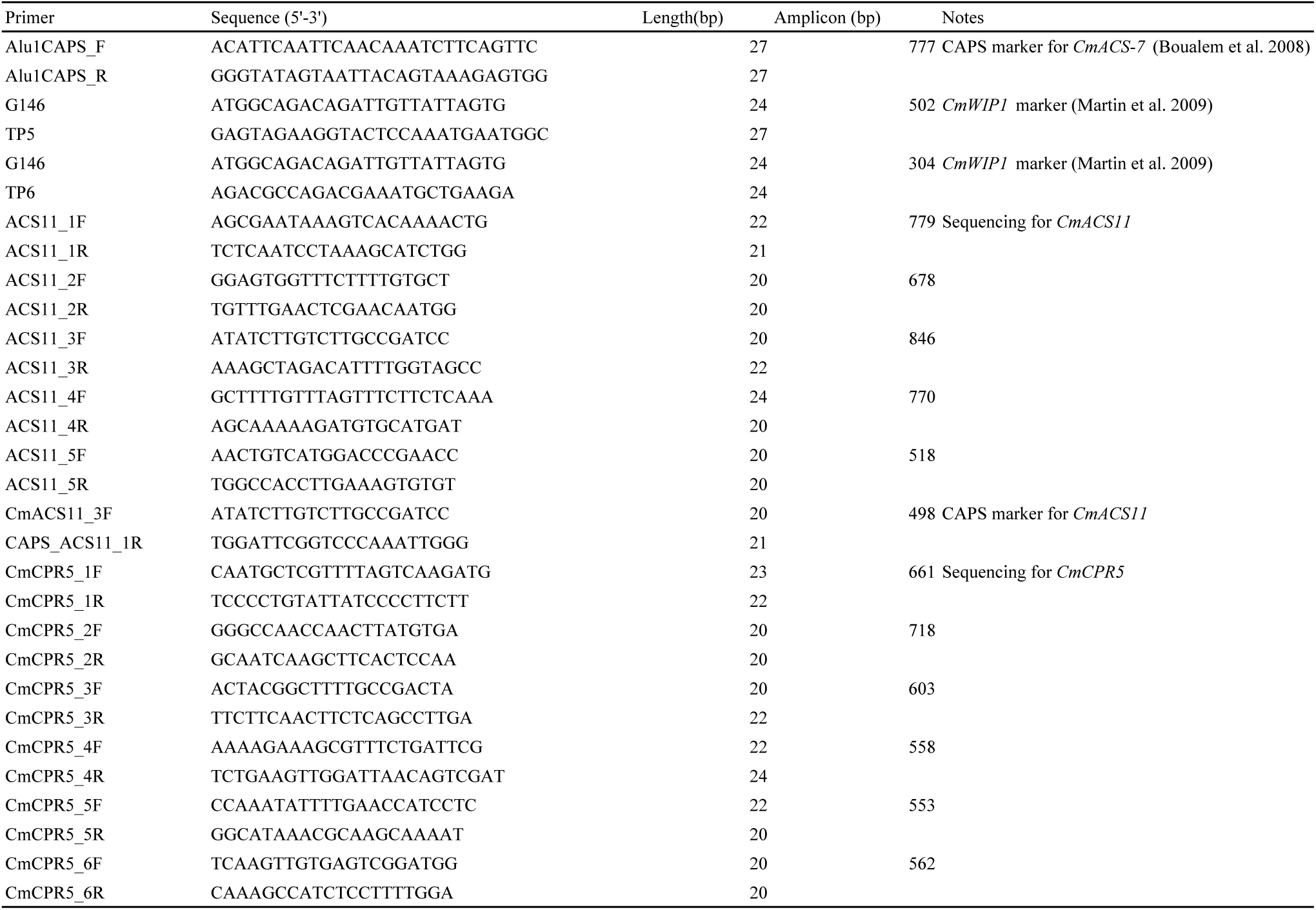
Primers for sequencing and CAPS analysis used in this study.

**Table S2.** Phenotypic variation for sex type on the main stem in the cultivation of 2015.

| Lines/Populations | No. of plants | Sex type on the main stem |  |  |
| --- | --- | --- | --- | --- |
|  |  | Femaleness | Maleness | Mixed |
| EF | 8 | 0 | 8 | 0 |
| UT1 | 8 | 8 | 0 | 0 |
| F<sub>1</sub> [EF (♀) × UT1 (♂)] | 8 | 8 | 0 | 0 |
| F<sub>1</sub> [UT1 (♀) × EF (♂)] | 8 | 8 | 0 | 0 |
| F<sub>2</sub> [EF (♀) × UT1 (♂)] | 136 | 89 | 26 | 21 |

**Table S3.** List of genes within the 1.5-LOD support interval of QTL for type of femeleness flower on Chr. 8 detected in the cultivation of 2020.

| Gene ID | Description (DHL v3.5.1) | Description (DHL v4) |
| --- | --- | --- |
| MELO3C007626 | Formin-like protein | Formin-like protein 6 |
| MELO3C007627 | BTB/POZ and TAZ domain protein | BTB/POZ and TAZ domain-containing protein 4 |
| MELO3C007628 | Cyclin family protein | B-like cyclin |
| MELO3C007629 | Condensin-2 complex subunit G2, putative | ARM repeat superfamily protein, putative isoform 1 |
| MELO3C007630 | GRAS family transcription factor | protein SHORT-ROOT |
| MELO3C007631 | Peptidyl-prolyl cis-trans isomerase-like 2 | RING-type E3 ubiquitin transferase |
| MELO3C007632 | Adenylyl-sulfate kinase | Adenylyl-sulfate kinase |
| MELO3C007633 | BTB/POZ domain-containing protein | No data |
| MELO3C007634 | MYB-family transcription factor, putative | transcription factor DIVARICATA-like |
| MELO3C007635 | 50S ribosomal protein L24 | 60S ribosomal protein L26 |
| MELO3C007636 | Calcium-transporting ATPase | Calcium-transporting ATPase |
| MELO3C007637 | Pentatricopeptide repeat-containing protein | No data |
| MELO3C007638 | Pentatricopeptide repeat-containing protein, putative | Pentatricopeptide repeat-containing protein |
| MELO3C007639 | 50S ribosomal protein L7/L12 | 50S ribosomal protein L7/L12 |
| MELO3C007640 | Putative MYB transcription factor | MYB transcription factor |
| MELO3C007877 | Lateral root primordium protein-related, putative | protein SHI RELATED SEQUENCE 1 |
| MELO3C007878 | Methyltransferase family protein, expressed | S-adenosyl-L-methionine-dependent methyltransferases superfamily protein |
| MELO3C007879 | Glutaredoxin-C3 | glutaredoxin-C3 |
| MELO3C007880 | Circadian clock protein KaiB | Oxidoreductase-like domain-containing protein |
| MELO3C007881 | Leucine-rich repeat receptor-like protein kinase family protein | Leucine-rich receptor-like protein kinase family protein |
| MELO3C007882 | GATA transcription factor 5, putative | GATA transcription factor |
| MELO3C007883 | Zinc finger CCCH domain-containing protein 30 | Zinc finger CCCH domain-containing protein 6-like |
| MELO3C007884 | Cytochrome P450 protein | cytochrome P450 84A1-like |
| MELO3C007885 | UPF0183 protein | UPF0183 protein At3g51130 |
| MELO3C007886 | B-box zinc finger family protein | Zinc finger, B-box |
| MELO3C007887 | Receptor-like protein kinase | Receptor-like protein kinase |
| MELO3C007641 | NADH dehydrogenase iron-sulfur protein 4 | NADH dehydrogenase [ubiquinone] iron-sulfur protein 4, mitochondrial |
| MELO3C007642 | Acetylglutamate kinase | Amino-acid N-acetyltransferase |
| MELO3C007643 | Amino-acid acetyltransferase | No data |
| MELO3C007644 | DUF4228 domain protein | Cyclin-D1-1-like |
| MELO3C007645 | Sodium channel protein type 8 subunit alpha | Unknown protein |
| MELO3C007646 | heptahelical protein 4 | heptahelical transmembrane protein 4-like |
| MELO3C007647 | Alpha galactosyltransferase family protein | galactomannan galactosyltransferase 1-like |
| MELO3C007648 | RuvB-like helicase 2 | RuvB-like helicase |
| MELO3C007649 | Phosphoribosylformylglycinamide cyclo-ligase | Uncharacterised conserved protein UCP031279 |
| MELO3C007650 | VQ motif-containing protein, putative | VQ domain-containing protein |
| MELO3C007651 | BSD domain-containing protein, putative | BSD domain-containing protein |
| MELO3C007652 | Bifunctional uridylyltransferase/uridylyl-removing enzyme | ACT domain-containing protein ACR1 |
| MELO3C007653 | Protein PHLOEM PROTEIN 2-LIKE A10 | Protein PHLOEM PROTEIN 2-LIKE A10-like |
| MELO3C007654 | Phytosulfokines, putative | Phytosulfokine-beta |
| MELO3C007655 | BZIP transcription factor | BZIP domain-containing protein |
| MELO3C007656 | Growth-regulating factor 1 | Growth-regulating factor |
| MELO3C007657 | AP2-like ethylene-responsive transcription factor | AP2-like ethylene-responsive transcription factor ANT |
| MELO3C007658 | Leucine-rich repeat receptor-like protein kinase | inactive LRR receptor-like serine/threonine-protein kinase BIR2 |
| MELO3C007659 | ATP-dependent RNA helicase | DEAD-box ATP-dependent RNA helicase |
| MELO3C007660 | DUF21 domain-containing-like protein | DUF21 domain-containing protein |
| MELO3C007661 | Coffea canephora DH200=94 genomic scaffold, scaffold_1 | Unknown protein |
| MELO3C007662 | 1-aminocyclopropane-1-carboxylate synthase | 1-aminocyclopropane-1-carboxylate synthase |
| MELO3C007663 | Myb transcription factor | transcription factor RAX2 |
| MELO3C007664 | Branched-chain-amino-acid aminotransferase | Branched-chain-amino-acid aminotransferase |
| MELO3C007665 | Putative nuclear matrix constituent protein 1-like protein | protein CROWDED NUCLEI 4 |
| MELO3C007666 | Homeobox-leucine zipper protein | Homeobox-leucine zipper protein |
| MELO3C007667 | Epidermal patterning factor-like protein 2 | Epidermal patterning factor-like protein |
| MELO3C007668 | Cysteine desulfurase IscS | Cysteine desulfurase |
| MELO3C007669 | Flagellar attachment zone protein 1 | No data |
| MELO3C007670 | Secretory carrier membrane protein (SCAMP) family protein | Unknown protein |
| MELO3C007671 | Coffea canephora DH200=94 genomic scaffold, scaffold_8 | No data |
| MELO3C007672 | Cys-Gly metalloprotease DUG1 | Cys-Gly metalloprotease DUG1 |
| MELO3C007673 | Tubulin alpha chain, putative | Tubulin alpha chain |
| MELO3C007674 | Malate dehydrogenase | Malate dehydrogenase |
| MELO3C007675 | Receptor-like protein kinase 3 | Leucine-rich receptor-like protein kinase family protein |
| MELO3C007676 | Receptor-like protein kinase 3 | No data |
| MELO3C007677 | Hexokinase | Phosphotransferase |
| MELO3C007678 | Basic helix loop helix (BHLH) family transcription factor | Basic helix-loop-helix transcription factor |
| MELO3C007679 | Basic helix-loop-helix DNA-binding superfamily protein, putative | transcription factor bHLH18-like |
| MELO3C007680 | basic helix-loop-helix (bHLH) DNA-binding superfamily protein | Basic helix-loop-helix (bHLH) DNA-binding family protein |
| MELO3C007681 | WD repeat-containing protein 5 | No data |
| MELO3C007682 | Kinesin-like protein | No data |
| MELO3C007683 | Kinesin-like protein | Kinesin-like protein |
| MELO3C007684 | Glycogen synthase | No data |
| MELO3C007685 | Glycogen synthase | glycogen synthase isoform X2 |
| MELO3C007686 | SPT2 chromatin protein | protein SPT2 homolog |
| MELO3C007687 | Phosphoenolpyruvate carboxykinase [ATP] | Phosphoenolpyruvate carboxykinase (ATP) |
| MELO3C007688 | tRNA-dihydrouridine synthase | tRNA-dihydrouridine synthase |
| MELO3C007689 | LisH/CRA/RING-U-box protein | protein RMD5 homolog |
| MELO3C007690 | Zinc/RING finger protein 4 | No data |
| MELO3C007691 | Auxin-responsive protein | Auxin-responsive protein |
| MELO3C007692 | Hydroxyproline-rich glycoprotein family protein | Hydroxyproline-rich glycoprotein family protein |
| MELO3C007693 | Putative homeodomain-like transcription factor superfamily protein | Homeodomain-like superfamily protein |
| MELO3C007694 | Transcription factor | transcription factor bHLH93 |
| MELO3C007695 | Putative beta-D-xylosidase 6 | Fn3_like domain-containing protein |
| MELO3C007696 | ABC transporter | ABC transporter G family member 17-like |
| MELO3C007697 | Protein BRICK1 | protein BRICK 1 |
| MELO3C007698 | Serine hydroxymethyltransferase | Serine hydroxymethyltransferase |
| MELO3C007699 | Coffea canephora DH200=94 genomic scaffold, scaffold_8 | Unknown protein |
| MELO3C007700 | Agamous MADS-box transcription factor | MADS-box transcription factor |
| MELO3C007701 | Rhamnogalacturonate lyase | Rhamnogalacturonan endolyase |
| MELO3C007702 | Alcohol dehydrogenase, putative | cinnamyl-alcohol dehydrogenase |
| MELO3C007703 | Elongator complex 4 | elongator complex protein 4 |
| MELO3C007704 | Dof zinc finger protein | dof zinc finger protein DOF5.7 |
| MELO3C007705 | Aldehyde dehydrogenase | Methylmalonate-semialdehyde dehydrogenase (CoA acylating) |
| MELO3C007706 | UDP-glycosyltransferase | UDP-glycosyltransferase 91A1-like |
| MELO3C007707 | UDP-glycosyltransferase | UDP-glycosyltransferase 91A1-like |
| MELO3C007708 | UDP-glycosyltransferase | UDP-glycosyltransferase 91A1-like |
| MELO3C007709 | Transcription initiation factor TFIID subunit 8, putative isoform 1 | BTP domain-containing protein |
| MELO3C007710 | Glycine--tRNA ligase beta subunit | No data |
| MELO3C007711 | Receptor protein kinase, putative | U-box domain-containing protein kinase family protein, putative |
| MELO3C007712 | Isochorismatase hydrolase family protein | nicotinamidase 1 |
| MELO3C007713 | Intermediate capsid protein VP6 | Forkhead box protein |
| MELO3C007714 | basic helix-loop-helix (bHLH) DNA-binding superfamily protein | ATP binding protein |
| MELO3C007715 | O-fucosyltransferase family protein | O-fucosyltransferase family protein |
| MELO3C007716 | MADS-box transcription factor | MADS-box protein SVP-like |
| MELO3C007717 | Kinesin-like protein | kinesin-like protein KCA1 |
| MELO3C007718 | GPI ethanolamine phosphate transferase 2 | GPI ethanolamine phosphate transferase 2 |
| MELO3C007719 | DnaJ domain containing protein, expressed | GPI ethanolamine phosphate transferase 2 |
| MELO3C007720 | Basic helix-loop-helix DNA-binding superfamily protein, putative | transcription factor bHLH131-like |
| MELO3C007721 | Glycine-rich protein | No data |
| MELO3C007722 | Impact-like protein | UPF0029 domain-containing protein |
| MELO3C007723 | Mitochondrial carrier protein | mitochondrial uncoupling protein 5-like |
| MELO3C007724 | DNA-directed RNA polymerase subunit beta' | Unknown protein |
| MELO3C007725 | Cyclin d4,2, putative | B-like cyclin |
| MELO3C007726 | Coffea canephora DH200=94 genomic scaffold, scaffold_7 | Cyclin N-terminal domain-containing protein |
| MELO3C007727 | Zinc finger homeodomain protein 1 | zinc-finger homeodomain protein 1-like |
| MELO3C007728 | GRAM domain protein/ABA-responsive-like protein | GRAM domain-containing protein |
| MELO3C007729 | Classical arabinogalactan protein 11 | vegetative cell wall protein gp1-like |
| MELO3C007730 | MATE efflux family protein | Protein DETOXIFICATION |
| MELO3C007731 | MATE efflux family protein | Protein DETOXIFICATION |
| MELO3C007732 | Ring finger protein, putative | RING-type domain-containing protein |
| MELO3C007733 | Protein of unknown function, DUF617 | protein MIZU-KUSSEI 1 |
| MELO3C007734 | Putative glucan 1,3-beta-glucosidase A | Mannan endo-1,4-beta-mannosidase |
| MELO3C007735 | Ring finger protein, putative | RING-type E3 ubiquitin transferase |
| MELO3C007736 | homeobox protein 6 | homeobox-leucine zipper protein ATHB-6-like |
| MELO3C007737 | LMBR1-like membrane protein | No data |
| MELO3C007738 | LMBR1 domain-containing protein 2 like A | LMBR1 domain-containing protein 2 homolog A |
| MELO3C007739 | Protein kinase 1 | serine/threonine-protein kinase SAPK2 |
| MELO3C007740 | 40S ribosomal protein S6 | 40S ribosomal protein S6 |
| MELO3C007741 | Ribosomal RNA small subunit methyltransferase B | tRNA (Cytosine(34)-C(5))-methyltransferase |
| MELO3C007742 | Putative Ras-related protein Rab | Ras-related protein Rab11A |
| MELO3C007743 | Polyadenylate-binding protein | Polyadenylate-binding protein |
| MELO3C007744 | 2-dehydro-3-deoxyphosphoheptonate aldolase (3-deoxy-d-arabino-heptulos | Phospho-2-dehydro-3-deoxyheptonate aldolase |
| MELO3C007745 | Transmembrane protein, putative | Transmembrane protein |
| MELO3C007746 | EPIDERMAL PATTERNING FACTOR-like protein 1 | Epidermal patterning factor-like protein |
| MELO3C007747 | Haloacid dehalogenase-like hydrolase | haloacid dehalogenase-like hydrolase domain-containing protein At4g39970 |
| MELO3C007748 | 1-deoxy-D-xylulose-5-phosphate synthase | Haloacid dehalogenase-like hydrolase domain-containing protein |
| MELO3C007749 | UDP-glycosyltransferase | No data |
| MELO3C007750 | Alpha/beta-hydrolase superfamily protein | Alpha/beta hydrolase-1 |
| MELO3C007751 | Coffea canephora DH200=94 genomic scaffold, scaffold_7 | Prostatic spermine-binding protein |
| MELO3C007752 | C2-H2 zinc finger protein | C2H2-type domain-containing protein |
| MELO3C007753 | MYB-related transcription factor | transcription factor MYB41-like |
| MELO3C007754 | Mitochondrial carrier protein, expressed | Mitochondrial carrier protein |
| MELO3C007755 | Ras-related protein Rab-6A | Rab family |
| MELO3C007756 | UDP-galactose transporter | CMP-sialic acid transporter 5 |
| MELO3C007757 | Asparagine synthetase | Asparagine synthetase [glutamine-hydrolyzing] |
| MELO3C007758 | Coffea canephora DH200=94 genomic scaffold, scaffold_7 | DUF4057 domain-containing protein |
| MELO3C007759 | C2 domain protein | C2 domain-containing protein |
| MELO3C007760 | Xyloglucan galactosyltransferase | Exostosin domain-containing protein |
| MELO3C007761 | Tetratricopeptide repeat 1 | Tetratricopeptide repeat protein 1 |
| MELO3C007762 | Mediator of RNA polymerase II transcription subunit 26, putative | Mediator of RNA polymerase II transcription subunit 26, putative |
| MELO3C007763 | Transporter-related family protein | MFS domain-containing protein |
| MELO3C007764 | DNA-directed RNA polymerase subunit beta | DUF1664 domain-containing protein |
| MELO3C007765 | VQ motif-containing protein | VQ domain-containing protein |
| MELO3C007766 | Protein SCO1 like 2, mitochondrial | protein SCO1 homolog 2, mitochondrial |
| MELO3C007767 | Lipase/lipoxygenase, PLAT/LH2 family protein | PLAT domain-containing protein |
| MELO3C007768 | Trehalose 6-phosphate phosphatase | Trehalose 6-phosphate phosphatase |
| MELO3C007769 | Ethylene-responsive transcription factor 5 | ethylene-responsive transcription factor ERF060 |
| MELO3C007770 | Acetylglutamate kinase | Unknown protein |
| MELO3C007771 | Type I inositol-1,4,5-trisphosphate 5-phosphatase | type I inositol polyphosphate 5-phosphatase 5-like |
| MELO3C007772 | 2,3-bisphosphoglycerate-dependent phosphoglycerate mutase | Phosphoglycerate mutase (2,3-diphosphoglycerate-dependent) |
| MELO3C007773 | 50s ribosomal L24e | FLZ-type domain-containing protein |
| MELO3C007774 | L-myo-inositol-1-phosphate synthase | Inositol-3-phosphate synthase |
| MELO3C007775 | DNA polymerase III polC-type | Exonuclease domain-containing protein |
| MELO3C007776 | Trafficking protein particle complex subunit 12 | trafficking protein particle complex subunit 12 |
| MELO3C007777 | Laccase | L-ascorbate oxidase |
| MELO3C007778 | Peptidyl-prolyl cis-trans isomerase | Peptidylprolyl isomerase |
| MELO3C007779 | Heavy metal transport/detoxification superfamily protein | heavy metal-associated isoprenylated plant protein 23 |
| MELO3C007780 | Eukaryotic aspartyl protease family protein | aspartic proteinase-like protein 1 isoform X1 |
| MELO3C007781 | U-box domain-containing 4-like protein | protein CELLULOSE SYNTHASE INTERACTIVE 1 |
| MELO3C007782 | MICOS complex subunit MIC60 | MICOS complex subunit MIC60 isoform X2 |
| MELO3C007783 | Similarity to dehydrogenase | NAD(P)-binding Rossmann-fold superfamily protein |
| MELO3C007784 | Coffea canephora DH200=94 genomic scaffold, scaffold_6 | Unknown protein |
| MELO3C007785 | RING-H2 finger protein 2B | E3 ubiquitin-protein ligase RHA2B-like |
| MELO3C007786 | DNA-directed RNA polymerase subunit beta | Unknown protein |
| MELO3C007787 | Zinc finger family protein | RING-CH-type domain-containing protein |
| MELO3C007788 | P-loop containing nucleoside triphosphate hydrolases superfamily protein | prostatic spermine-binding protein isoform X2 |
| MELO3C007789 | Gamma-glutamyltranspeptidase 1 | glutathione hydrolase 1 |
| MELO3C007790 | UvrABC system protein C | Translation initiation factor IF-2, putative isoform 1 |
| MELO3C007791 | Pentatricopeptide repeat-containing protein, putative | Pentatricopeptide repeat-containing protein |
| MELO3C007792 | Cytochrome P450 family protein | cytochrome P450 CYP736A12-like |
| MELO3C007793 | Cytochrome P450 family protein | cytochrome P450 CYP736A12-like |
| MELO3C007794 | Cytochrome P450 family protein | cytochrome P450 CYP736A12-like |
| MELO3C007795 | Cytochrome P450 family protein | cytochrome P450 CYP736A12-like |
| MELO3C007796 | Cytochrome P450 | Cytochrome P450 CYP736A12-like protein |
| MELO3C007797 | Cytochrome P450 family protein | cytochrome P450 CYP736A12-like |
| MELO3C007798 | Cytochrome P450 family protein | Cytochrome P450 family protein |
| MELO3C007799 | Cytochrome P450 family protein | cytochrome P450 CYP736A12-like |
| MELO3C007800 | Receptor protein kinase, putative | Protein kinase domain-containing protein |
| MELO3C007801 | Probable protein kinase UbiB | protein ACTIVITY OF BC1 COMPLEX KINASE 8, chloroplastic-like |
| MELO3C007802 | CPR5, putative | protein CPR-5-like |
| MELO3C007803 | CPR5, putative | protein CPR-5-like |
| MELO3C007804 | BRASSINAZOLE-RESISTANT 1 protein, putative | BES1/BZR1 homolog protein 2 |
| MELO3C007805 | UDP-glycosyltransferase 1 | UDP-glycosyltransferase 72E1 |
| MELO3C007806 | Putative glycerophosphoryl diester phosphodiesterase 3 | Glycerophosphodiester phosphodiesterase |
| MELO3C007807 | Xaa-pro aminopeptidase, putative | aminopeptidase P1 |
| MELO3C007808 | NAD(P)H dehydrogenase (Quinone) | NAD(P)H dehydrogenase (quinone) |
| MELO3C007809 | homeobox protein 21 | homeobox-leucine zipper protein ATHB-40 |
| MELO3C007810 | Protein kinase superfamily protein | Protein kinase domain-containing protein |
| MELO3C007811 | Transcription factor | G-box-binding factor 1-like |
| MELO3C007812 | HVA22-like protein | HVA22-like protein |
| MELO3C007813 | Translation initiation factor IF-2 | vicilin-like seed storage protein At2g18540 |
| MELO3C007814 | Zinc finger protein | protein indeterminate-domain 2-like |
| MELO3C007815 | Putative polyol transporter 6 | Major facilitator superfamily protein |
| MELO3C007816 | Nonsense-mediated mRNA decay NMD3 family protein | 60S ribosomal export protein NMD3 |
| MELO3C007817 | Isoleucine--tRNA ligase | Protein of unknown function (DUF620) |
| MELO3C007818 | Sugar transporter, putative | Protein of unknown function (DUF1195) |
| MELO3C007819 | Transcription initiation factor IIB | Plant-specific TFIIB-related protein 1 |
| MELO3C007820 | Anti-Muellerian hormone type-2 receptor | Anti-muellerian hormone type-2 receptor |
| MELO3C007821 | RNA binding protein, putative | U1 small nuclear ribonucleoprotein 70 kDa |
| MELO3C007822 | Zinc finger and BTB domain-containing protein 22 | Unknown protein |
| MELO3C007823 | Scarecrow-like transcription factor 7 family protein | Scarecrow-like protein 4 |
| MELO3C007824 | Late embryogenesis abundant protein | Late embryogenesis abundant protein |
| MELO3C007825 | Phosphatidylinositol transfer protein | CRAL-TRIO domain-containing protein |
| MELO3C007826 | Methyltransferase family protein, putative | protein N-lysine methyltransferase METTL21A |
| MELO3C007827 | Soluble pyridine nucleotide transhydrogenase | pyridine nucleotide-disulfide oxidoreductase domain-containing protein 2-like |
| MELO3C007828 | 50S ribosomal protein L6, putative | 50S ribosomal protein L6, putative |
| MELO3C007829 | Putative ADP-ribosylation factor | ADP-ribosylation factor family protein |
| MELO3C007830 | DNA-3-methyladenine glycosylase 1 | DNA-3-methyladenine glycosylase 1-like |
| MELO3C007831 | GATA transcription factor, putative | GATA transcription factor 18 |
| MELO3C007832 | Zinc finger-homeodomain protein 2 | Zinc-finger homeodomain protein 2 |
| MELO3C007833 | AAA-ATPase-like protein | ATPase family AAA domain-containing protein 3 |
| MELO3C007834 | Tubby-like F-box protein | Tubby-like F-box protein |
| MELO3C007835 | Pentatricopeptide repeat | Pentatricopeptide repeat-containing protein |
| MELO3C007836 | Transmembrane-like protein | transmembrane protein 53 |
| MELO3C007837 | Replication factor C subunit 3 | replication factor C subunit 5-like |
| MELO3C007838 | Inorganic pyrophosphatase, putative | Inorganic diphosphatase |
| MELO3C007839 | Chaperone DnaJ-domain protein, putative | auxilin-like protein 1 isoform X1 |
| MELO3C007840 | 3-hydroxyacyl-[acyl-carrier-protein] dehydratase FabZ | Auxilin-like protein 1 isoform X4 |
| MELO3C007841 | 1,4-alpha-glucan branching enzyme GlgB | Unknown protein |
| MELO3C007842 | Mannan endo-1,4-beta-mannosidase 7 | Mannan endo-1,4-beta-mannosidase |
| MELO3C007843 | Phosphatidic acid phosphatase family protein, putative | lipid phosphate phosphatase epsilon 1, chloroplastic-like |
| MELO3C007844 | Coffea canephora DH200=94 genomic scaffold, scaffold_7 | Unknown protein |
| MELO3C007845 | Methyltransferase-like 3 | S-adenosyl-L-methionine-dependent methyltransferases superfamily protein |
| MELO3C007846 | Methyltransferase-like 2 | S-adenosyl-L-methionine-dependent methyltransferases superfamily protein |
| MELO3C007847 | Methyltransferase-like 2 | S-adenosyl-L-methionine-dependent methyltransferases superfamily protein |
| MELO3C007848 | Putative N-methyltransferase | S-adenosyl-L-methionine-dependent methyltransferases superfamily protein |
| MELO3C007849 | N-methyltransferase | S-adenosyl-L-methionine-dependent methyltransferases superfamily protein |
| MELO3C007850 | Methyltransferase-like 2 | S-adenosyl-L-methionine-dependent methyltransferases superfamily protein |
| MELO3C007851 | Methyltransferase | No data |
| MELO3C007852 | Carboxyl methyltransferase 4 | S-adenosyl-L-methionine-dependent methyltransferases superfamily protein |
| MELO3C007853 | Heavy metal transport/detoxification superfamily protein, putative | heavy metal-associated isoprenylated plant protein 3-like |
| MELO3C007854 | UPF0261 protein | UPF0261 protein |
| MELO3C007855 | Pre-mrna-splicing factor | Protein pleiotropic regulatory locus 1 |
| MELO3C007856 | Cytochrome P450 family protein | abscisic acid 8'-hydroxylase 1-like |
| MELO3C007857 | ATP-dependent zinc metalloprotease FTSH 8, mitochondrial | P-loop containing nucleoside triphosphate hydrolases superfamily protein |
| MELO3C007858 | ATP-dependent zinc metalloprotease FTSH 8, mitochondrial | No data |
| MELO3C007859 | ATP-dependent zinc metalloprotease FTSH 4, mitochondrial | AAA-ATPase |
| MELO3C007860 | ATP-dependent zinc metalloprotease FTSH 10, mitochondrial | AAA-ATPase At2g18193-like |
| MELO3C007861 | P-loop containing nucleoside triphosphate hydrolases superfamily protein | AAA-ATPase At2g18193-like |
| MELO3C007862 | ATP-dependent zinc metalloprotease FTSH 4, mitochondrial | AAA-ATPase |
| MELO3C007863 | Sec14p-like phosphatidylinositol transfer family protein | phosphatidylinositol/phosphatidylcholine transfer protein SFH12 |
| MELO3C007864 | Pentatricopeptide repeat (PPR) superfamily protein | basic helix-loop-helix (bHLH) DNA-binding superfamily protein |
| MELO3C007865 | Transcription factor | bZIP transcription factor 53-like |
| MELO3C007866 | CONTAINS InterPro DOMAIN/s: Ubiquitin-conjugating enzyme E2C-binc | Ubiquitin-conjugating enzyme E2-binding protein |
| MELO3C007867 | Putative uncharacterized protein At4g36440 | Unknown protein |
| MELO3C007868 | Peroxidase | No data |
| MELO3C007869 | 50S ribosomal protein L7/L12 | 50S ribosomal protein L7/L12 |
| MELO3C007870 | Cytochrome P450 superfamily protein | 3-epi-6-deoxocathasterone 23-monooxygenase |
| MELO3C007871 | Ribosomal RNA small subunit methyltransferase A | rRNA adenine N(6)-methyltransferase |
| MELO3C007872 | Beta-galactosidase | Beta-galactosidase |
| MELO3C007873 | Thioredoxin | thioredoxin H-type |
| MELO3C007874 | Purple acid phosphatase | Purple acid phosphatase |
| MELO3C007875 | protein phosphatase 2A-4 | Unknown protein |
| MELO3C007876 | FG-GAP repeat-containing protein | FG-GAP repeat-containing protein |

**Table S4.** List of genes within the 1.5-LOD support interval of QTL for the femaleness on the main stem on Chr. 3 detected in the cultivation of 2020.

| Gene ID | Description (DHL v3.5.1) | Description (DHL v4) |
| --- | --- | --- |
| MELO3C010800 | F-box protein | F-box protein |
| MELO3C010799 | Exocyst subunit exo70 family protein G1 | Exocyst subunit Exo70 family protein |
| MELO3C010798 | NADH dehydrogenase iron-sulfur protein 4 | NADH dehydrogenase [ubiquinone] iron-sulfur protein 4, mitochondrial |
| MELO3C010797 | NACHT, LRR and PYD domains-containing protein 5 | glutamic acid-rich protein |
| MELO3C010796 | Coffea canephora DH200=94 genomic scaffold, scaffold_6 | Protein SOB FIVE-LIKE 5 |
| MELO3C010795 | Calcium-transporting ATPase | Calcium-transporting ATPase |
| MELO3C010794 | Calcium-transporting ATPase | No Data |
| MELO3C010792 | DHFS-FPGS homolog D | Unknown protein |
| MELO3C010791 | U-box domain-containing protein 30 | RING-type E3 ubiquitin transferase |
| MELO3C010790 | V-type proton ATPase subunit G3 | V-type proton ATPase subunit G |
| MELO3C010789 | CW7 protein, putative | CW7 protein, putative |
| MELO3C010787 | Kinesin-like protein | Kinesin-like protein |
| MELO3C010786 | Growth-regulating factor 1 | Growth-regulating factor |
| MELO3C010785 | Glutathione transport system permease protein GsiD | Unknown protein |
| MELO3C010784 | AP2-like ethylene-responsive transcription factor | AP2-like ethylene-responsive transcription factor ANT |
| MELO3C010783 | Transcription factor | transcription factor bHLH49 isoform X1 |
| MELO3C010782 | Cellulose synthase-like C1-1, glycosyltransferase family 2 protein | Cellulose synthase-like protein |
| MELO3C010781 | Squalene monooxygenase | Squalene monooxygenase |
| MELO3C010780 | Chaperone protein DnaJ | DnaJ domain containing protein |
| MELO3C010779 | 1-aminocyclopropane-1-carboxylate synthase | 1-aminocyclopropane-1-carboxylate synthase |
| MELO3C010778 | ERD (Early-responsive to dehydration stress) family protein | CSC1-like protein RXW8 isoform X1 |
| MELO3C010777 | Esterase/lipase/thioesterase family protein | transcription factor RAX3-like |
| MELO3C010776 | Branched-chain-amino-acid aminotransferase | Branched-chain-amino-acid aminotransferase |
| MELO3C010775 | Homeobox transcription factor KN4 | homeobox protein knotted-1-like 1 |
| MELO3C010774 | Zinc finger CCCH domain-containing protein 49 | Zinc finger CCCH domain-containing protein 20 |
| MELO3C010773 | Myb transcription factor | transcription factor RAX3-like |
| MELO3C010772 | Clathrin interactor EPSIN 1-like protein | Clathrin interactor EPSIN 1 |
| MELO3C010771 | Exonuclease 3'-5' domain-containing 1 | piRNA biogenesis protein EXD1 isoform X1 |
| MELO3C010769 | EARLY FLOWERING 3 | protein EARLY FLOWERING 3 |
| MELO3C010770 | alpha/beta-Hydrolases superfamily protein | Unknown protein |
| MELO3C010768 | Protein of Unknown Function (DUF239) | Protein of Unknown Function (DUF239) |
| MELO3C010767 | Plant invertase/pectin methyltransferase inhibitor superfamily protein | Unknown protein |
| MELO3C010766 | UPF0587 protein C1orf123-like protein | UPF0587 protein C1orf123 homolog |
| MELO3C010765 | arabinogalactan protein 15 | Unknown protein |
| MELO3C010764 | Protein of unknown function (DUF1639) | Protein of unknown function (DUF1639) |
| MELO3C010763 | Vacuolar processing enzyme | vacuolar-processing enzyme-like |
| MELO3C010762 | Ferrochelatase | Ferrochelatase |
| MELO3C010761 | RNA-binding KH domain protein | Far upstream element-binding protein 1 |
| MELO3C010760 | Pollen specific protein sf21 | pollen-specific protein SF21 isoform X1 |
| MELO3C010759 | BEL1-like homeodomain protein | homeobox protein ATH1 |
| MELO3C010758 | Myb-like transcription factor family protein, putative | myb-like transcription factor family protein |
| MELO3C010757 | Glutathione-regulated potassium-efflux system protein | K(+) efflux antiporter 6 |
| MELO3C010756 | Calcineurin B-like protein 10 | calcineurin B-like protein 10 |
| MELO3C010755 | At5g11810 | Recoverin |
| MELO3C010754 | Glycine dehydrogenase decarboxylating protein | Glycine cleavage system P protein |
| MELO3C010753 | Zinc transporter 4 | Zinc transporter 4 |
| MELO3C010752 | LOB domain-containing 22-like protein | LOB domain-containing protein |
| MELO3C010751 | Beta-fructofuranosidase; cell wall invertase I; fructosidase | Beta-fructofuranosidase, insoluble isoenzyme CWINV3-like |
| MELO3C010750 | Mitochondrial carrier family | ADP/ATP translocase |
| MELO3C010749 | Glutaredoxin | Glutaredoxin |
| MELO3C010748 | Heat shock transcription factor A2 | Heat stress transcription factor |
| MELO3C010747 | Calmodulin-binding family protein | IQ domain-containing protein IQM1 |
| MELO3C010746 | Mitochondrial carrier protein-like | mitochondrial carrier protein CoAc1-like |
| MELO3C010745 | Mitochondrial import receptor subunit TOM20-2 family protein | mitochondrial import receptor subunit TOM20-like |
| MELO3C010744 | CTD small phosphatase-like protein 2 | CTD small phosphatase-like protein 2 |
| MELO3C010743 | HORMA domain-containing 1 | meiosis-specific protein ASY1 |
| MELO3C010742 | MYND type zinc finger protein | F-box protein At1g67340-like |
| MELO3C010741 | Chaperone protein dnaJ-like protein | Heat shock protein DnaJ, cysteine-rich domain containing protein |
| MELO3C010740 | Glutamyl-tRNA reductase | Unknown protein |
| MELO3C010739 | Acyl-[acyl-carrier-protein] desaturase | Acyl-[acyl-carrier-protein] desaturase |
| MELO3C010738 | Thiosulfate sulfurtransferase GlpE | Rhodanese domain-containing protein |
| MELO3C010737 | U-box domain-containing protein 6 | RING-type E3 ubiquitin transferase |
| MELO3C010736 | KH domain-containing protein | Gelsolin-related protein of 125 kDa isoform X1 |
| MELO3C010735 | Nucleolar GTP-binding protein 1 | Nucleolar GTP-binding protein 1 |
| MELO3C010734 | ATP-dependent RNA helicase, putative | RNA helicase family protein |
| MELO3C010733 | ATP-dependent RNA helicase, putative | No Data |
| MELO3C010732 | Extracellular ligand-gated ion channel protein | Protein of unknown function (DUF3537) |
| MELO3C010731 | Kinase family protein | cyclin-dependent kinase G-2-like |
| MELO3C010730 | Remorin family protein | Remorin family protein |
| MELO3C010729 | Coffea canephora DH200=94 genomic scaffold, scaffold_74 | Serine/arginine repetitive matrix protein 2 |
| MELO3C010728 | Protein Iojap/ribosomal silencing factor RsfS | protein Iojap-related, mitochondrial |
| MELO3C010727 | Dehydrogenase/reductase SDR family member 4 | 3-oxoacyl-[acyl-carrier-protein] reductase |
| MELO3C010726 | Ankyrin repeat-containing protein | Ankyrin repeat-containing protein |
| MELO3C010725 | B12D protein | NADH-ubiquinone reductase complex 1 MLRQ subunit |
| MELO3C010724 | Trihelix transcription factor | trihelix transcription factor GT-1-like isoform X2 |
| MELO3C010723 | Restriction endonuclease, type II-like protein | YqaJ domain-containing protein |

**Table S5.** Genotypes of two sex determination genes and the SNP markers adjacent to *CmCPR5*, and sexual type segragation in F_2_ population in the cultivation of 2020.

| Genotype<sup>1</sup> | Expected phenotype<sup>2</sup> | Allele on the SNP marker<sup>3</sup> | Sexual type |  |  |
| --- | --- | --- | --- | --- | --- |
|  |  |  | Female | Bisexual | Both |
| *GGMM* | Female | UT1 | 0 | 7 | 0 |
|  |  | EF | 7 | 0 | 0 |
|  |  | Heterozygous | 3 | 6 | 1 |
| *GGMm* | Female | UT1 | 0 | 9 | 1 |
|  |  | EF | 16 | 1 | 4 |
|  |  | Heterozygous | 2 | 29 | 7 |
|  |  | No data | 1 | 2 | 0 |
| *GGmm* | Bisexual | UT1 | 0 | 5 | 1 |
|  |  | EF | 0 | 4 | 0 |
|  |  | Heterozygous | 0 | 12 | 2 |

<sup>1</sup> *G/g* : *CmWIP1* , *M/m* : *CmACS-7*
<sup>2</sup> Expected phenotype based on the genotypes in *CmWIP1* and *CmACS-7*
<sup>3</sup> Based on SNP marker S8\_5329551, approximately 30,000 bp away from *CmCPR5*

**Table S6.** Frequency of femaleness, maleness and mixed phenotypes on the main stem in F_2_ population derived from a cross between a non-netted andromonoecious fixed line (HOF) and UT1.

| Genotype | No. of plants | Femaleness | Maleness | Mixed |
| --- | --- | --- | --- | --- |
| UT1 allele | 17 | 15 | 0 | 2 |
| Hetero | 30 | 20 | 2 | 8 |
| HOF allele | 13 | 0 | 13 | 0 |
| Total | 60 | 35 | 15 | 10 |

